# Identification of the *Staphylococcus aureus* Endothelial Cell Surface Interactome by Proximity Labeling

**DOI:** 10.1101/2024.11.26.625535

**Authors:** Marcel Rühling, Fabio Schmelz, Alicia Kempf, Kerstin Paprotka, Martin J. Fraunholz

## Abstract

Virulence strategies of pathogens depend on interaction with host cells. Binding and activation of receptors located on the plasma membrane is crucial for the attachment to or pathogen internalization by host cells. Identification of host cell receptors often is difficult and hence the identity of many proteins that play important roles during host-pathogen interaction remains elusive. We developed a novel proximity labeling approach, in which we decorated the opportunistic pathogen *Staphylococcus aureus* with ascorbate peroxidase 2. Upon addition of hydrogen peroxide the peroxidase initiates proximity biotinylation of *S. aureus* host receptors, thereby enabling identification of these proteins. We here demonstrate an endothelial cell surface interactome of 306 proteins of which neuronal adhesion molecule, protein tyrosine kinase PTK7, melanotransferrin, protein-tyrosine kinase Met, CD109 and others constitute novel *S. aureus* co-receptors. Filtering the interactome for validated surface proteins resulted in a list of 89 receptor candidates of which a 53% were described to interact with *S. aureus* or other pathogens.

## Introduction

Pathogens interact with host cell surfaces through a variety of mechanisms that facilitate their entry, survival, and replication within the host. These interactions are crucial for the establishment of infections and can vary significantly among different types of pathogens, including bacteria [1], viruses [2] and protozoa [3]. Thereby, pathogens first need to adhere to host cells, which is often mediated through specific interactions with cell surface receptors. For example, many bacteria possess adhesins - surface proteins that bind to host cell receptors, allowing them to colonize, e.g., epithelial surfaces [1]. Adherence is a prerequisite for pathogen entry into host cells. Bacteria mainly are internalized either actively inducing their uptake, for instance, by injection of effectors via specialized secretion systems, termed trigger-type phagocytosis, or by the zipper-type mechanism, which is mediated by receptor-driven actin polymerization [4]. Intracellular localization has beneficial effects for the bacteria such as immune evasion or enhanced resistance to antibiotics [5-8]. Intensive research has enabled the identification of many receptors over the last decades [1, 9-12]. Nevertheless, reliable, and efficient methods for identifying host-pathogen interaction partners in their entirety are lacking.

*Staphylococcus aureus* is an opportunistic facultative intracellular pathogen that causes diseases ranging from soft tissue and skin infections to sepsis and toxic shock [13-15]. We know today that *S. aureus* possesses intracellular virulence and survival strategies in either phagocytes or even non-phagocytic cells that readily internalize *S. aureus* [16, 17]. Invasion of the latter, for example epithelial or endothelial cells, is mediated by α_5_β_1_ integrins on host cells that bind via fibronectin to fibronectin-binding proteins, which are covalently anchored in the staphylococcal cell wall [10]. To date, several entry receptors for staphylococcal invasion have been described, indicating a complex interface between pathogen and host cells [18-25].

We recently described that *S. aureus* can be internalized by host cells via a rapid pathway that takes place within minutes after the bacteria contact the host cell plasma membrane. This unique pathway depends on host cell Ca^2+^ signaling as well as the lysosomal enzyme acid sphingomyelinase (ASM). We hypothesized that the rapid internalization of *S. aureus* by host cells required the interaction with a specific subset of host proteins [26].

Identification of host receptors for pathogens is challenging. Genetic approaches, which frequently are based on RNAi or CRISPR technologies, enable high throughput screening for infection relevant proteins [27]. However, these methods are limited to receptors that measurably reduce the cellular uptake of the pathogen. Moreover, a genetic knock-out may permanently alter the composition of cell surface signaling complexes and may not necessarily constitute direct interaction partners of the receptor complex in question. Cross-linking approaches more reliably identify direct host-pathogen interaction partners. Thereby, artificial cross linkers are used to couple proteins on the pathogen surface with host cell receptors [28]. However, interaction partners that do not directly bind to the pathogen, but may serve important functions during infection, likely are missed.

We here present a novel, alternative technique for the identification of host cell receptors in direct proximity to the adhered pathogen, which is based on proximity biotinylation using ascorbate peroxidase 2 (APEX2). APEX2 is an enzyme, which allows for proximity biotinylation with high temporal resolution [29] enabling the identification of interaction partners with shorter labelling pulses, when compared to other techniques such as BioID or TurboID [30, 31]. Using this technique we here identified and validate neuronal adhesion molecule (NRCAM), protein tyrosine kinase PTK7, melanotransferrin (MELTF), the protein-tyrosine kinase Met and CD109 as novel *S. aureus* co-receptors.

## Results

### Decorating *S. aureus* with APEX2

To identify novel host cell receptors interacting with *S. aureus*, we designed an ascorbate peroxidase 2 (APEX2)-based proximity labeling approach (**Figure 1**, A), whereby we decorated *S. aureus* with a fusion protein consisting of APEX2, the fluorescent marker yellow-fluorescent protein (YFP) and the cell wall-targeting domain of the staphylolytic protease lysostaphin (CWT), which binds to the staphylococcal cell wall with high affinity [32]. The construct was cytosolically produced in HeLa cells, isolated, and subsequently used to decorate the bacteria. Efficient attachment of APEX2 to the staphylococcal surface was validated by flow cytometry, where we observed a higher YFP mean fluorescence of bacteria treated with cytosol containing APEX2-YFP-CWT (**Figure 1**, B). Next, we performed infection of unmodified HeLa cells with decorated bacteria in presence of biotin phenol. Proximity biotinylation was initiated by addition of H_2_O_2_, the reaction was stopped with quencher solution and samples were fixed. We stained biotin with streptavidin-PE and *S. aureus* with an anti-*S. aureus* antibody, while we used a fixable ceramide analog [33] to visualize host cells (**Figure 1**, C). As expected, biotin was detected in close vicinity to the bacteria, indicating that bacteria-bound APEX2 is functional and enables proximity biotinylation of host cell surface proteins.

**Figure 1.**
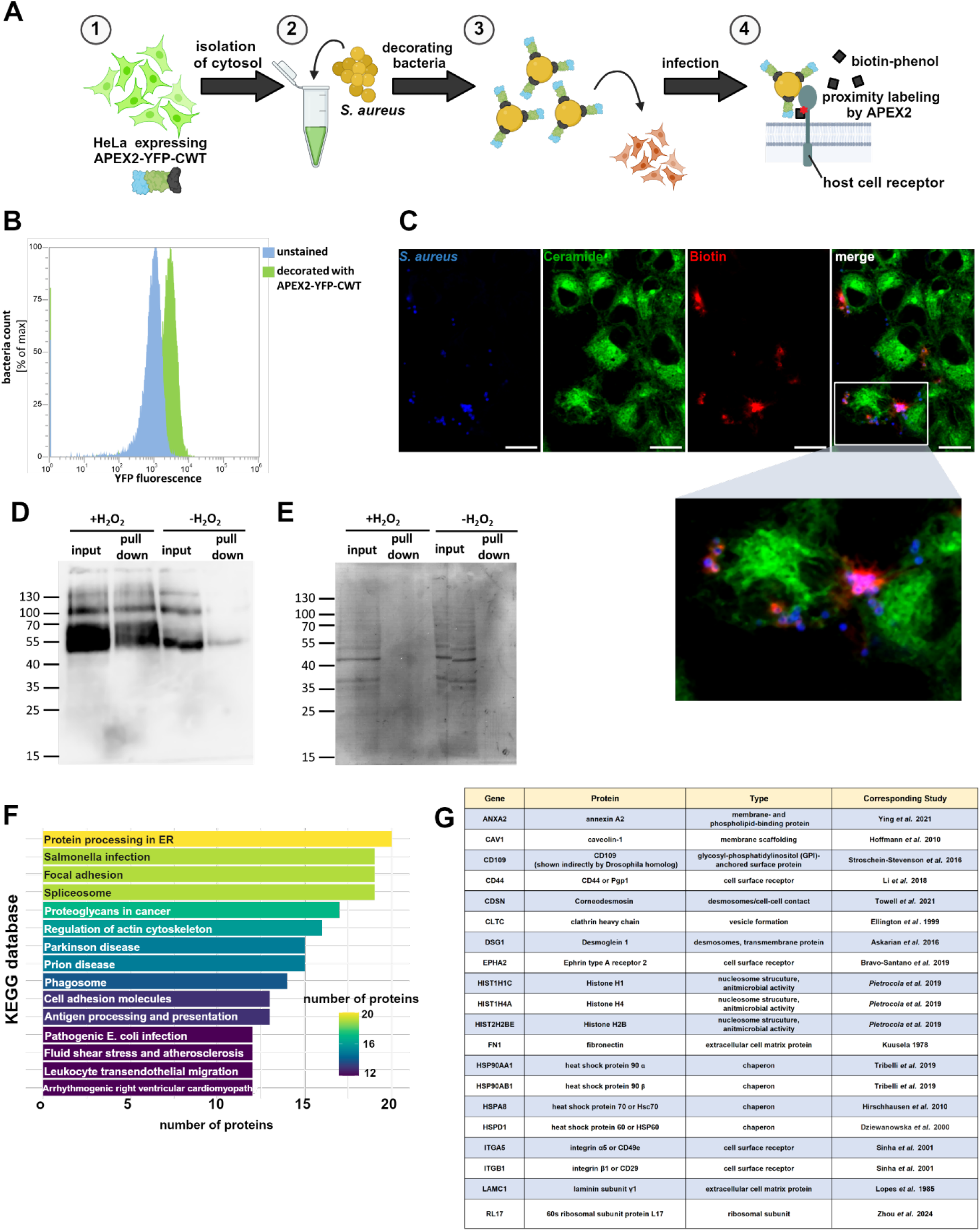
APEX2-based proximity labeling enables the detection of *S. aureus*-host interactome. (A) The cytosol of HeLa expressing the reporter APEX2-YFP-CWT was isolated (1) and used to decorate *S. aureus* JE2 *in vitro* (2). HuLEC were infected with the bacteria for 15 min (3) and proximity labeling of host receptors was initiated by addition of biotin phenol and a 1 min pulse of H_2_O_2_ (4). (**B) APEX2-YFP-CWT is attached to the *S. aureus* surface**. *S. aureus* was treated with a cytosol preparation of the APEX2-YFP-CWT cell line or left untreated. Bacteria were analyzed for YFP fluorescence via flow cytometry. **(C) APEX2-decoration of S. aureus enables local biotinylation**. HeLa were infected with APEX2-decorated bacteria in presence of biotin phenol. Biotinylation was induced by a 1 min H_2_O_2_ pulse, samples were fixed and stained for biotin via streptavidin-PE, α-NH_2_-ω-N_3_-C_6_-ceramide via BODIPY-Fl-DBCO and *S. aureus* via an anti-*S. aureus* antibody. Scale bars: 15 µm. (**D, E) Host proteins are biotinylated by APEX2-decorated bacteria and can be isolated by streptavidin pull down**. HuLEC were infected with APEX2-decorated bacteria in presence of biotin phenol. Biotinylation was induced by a 1 min H_2_O_2_ pulse, or samples were left untreated. Cells were lysed and biotinylated proteins were isolated by streptavidin pull down. Lysates and pull-down fractions were analyzed by anti-biotin Western blot (**C**) or Ponceau S staining (**D**). (**F) *S. aureus*-host interaction partners are associated with infection-related pathways**. Proteins that were identified by APEX2-based proximity labeling (n=3) were analyzed by pathfindR and categorized with the KEGG database. The number of gene associating with the most abundant categories are shown. **(G) APEX2 proximity labeling identifies known *S. aureus* interacting proteins**. The *S. aureus* host interactome identified by proximity labeling was searched for proteins that already are known for interacting with *S. aureus* [10, 18, 19, 21-25, 34-40].

In order to identify potential *S. aureus* receptors on the surface of human lung microvascular endothelial cells (HuLEC), we infected the cells with APEX2-decorated *S. aureus*, initiated proximity biotinylation with H_2_O_2_ and isolated biotinylated proteins by streptavidin pull down. We detected significantly more biotinylated proteins in pull down fractions, when H_2_O_2_ was added during infection when compared to the untreated control (**Figure 1**, D, E).

### Proximity labeling identifies NRCAM, PTK7, MELTF, CD109 and MET as ASM-dependent coreceptors of *S. aureus*

In total, mass spectrometry of the pull-down fractions identified 306 potential *S. aureus* interaction partners in three biological replicates (**Extended Data 1**). Pathway analysis identified “*Salmonella* infection” as well as “focal adhesions” as most enriched terms (**Figure 1**, F), indicating that infection relevant proteins were detected with our approach, since focal adhesions are well-known cellular entry sites for *S. aureus* [10]. StringDB detected a highly connected protein network [protein-protein interaction (PPI) enrichment p-value <1×10^−16^] and the accumulation of several infection-related proteins (**Supp. Figure 1**).

Of the 305 identified proteins, at least 20 were previously reported to interact with *S. aureus*, thereby corroborating our methodology (**Figure 1**, G).

Recently, we described that *S. aureus* is internalized by host cells via a rapid pathway that depends on host cell Ca^2+^ signaling and the lysosomal enzyme ASM [26]. This rapid pathway predominantly mediates host cell entry early during infection. We hypothesized that rapid internalization of *S. aureus* by host cells required the interaction with a specific subset of host molecules, which we set out to identify with the proximity labeling approach.

Therefore, we blocked the ASM-dependent uptake by the ASM inhibitor amitriptyline, the Ca^2+^ ionophore ionomycin or by removal of the ASM substrate sphingomyelin from the plasma membrane by treatment with the bacterial sphingomyelinase β-toxin. We next infected the cells with APEX2-decorated bacteria. Despite their APEX2 decoration the bacteria were still efficiently internalized by host cells, and invasion was sensitive to inhibitor treatments as observed for undecorated bacteria (**Figure 2**, A). Further, the removal of sphingomyelin had no effect on bacterial adherence to host cells (**Supp. Figure 1, A**), excluding that changes in biotinylation intensity arise from lower numbers of bacteria that adhered to host cells in treated samples.

**Figure 2.**
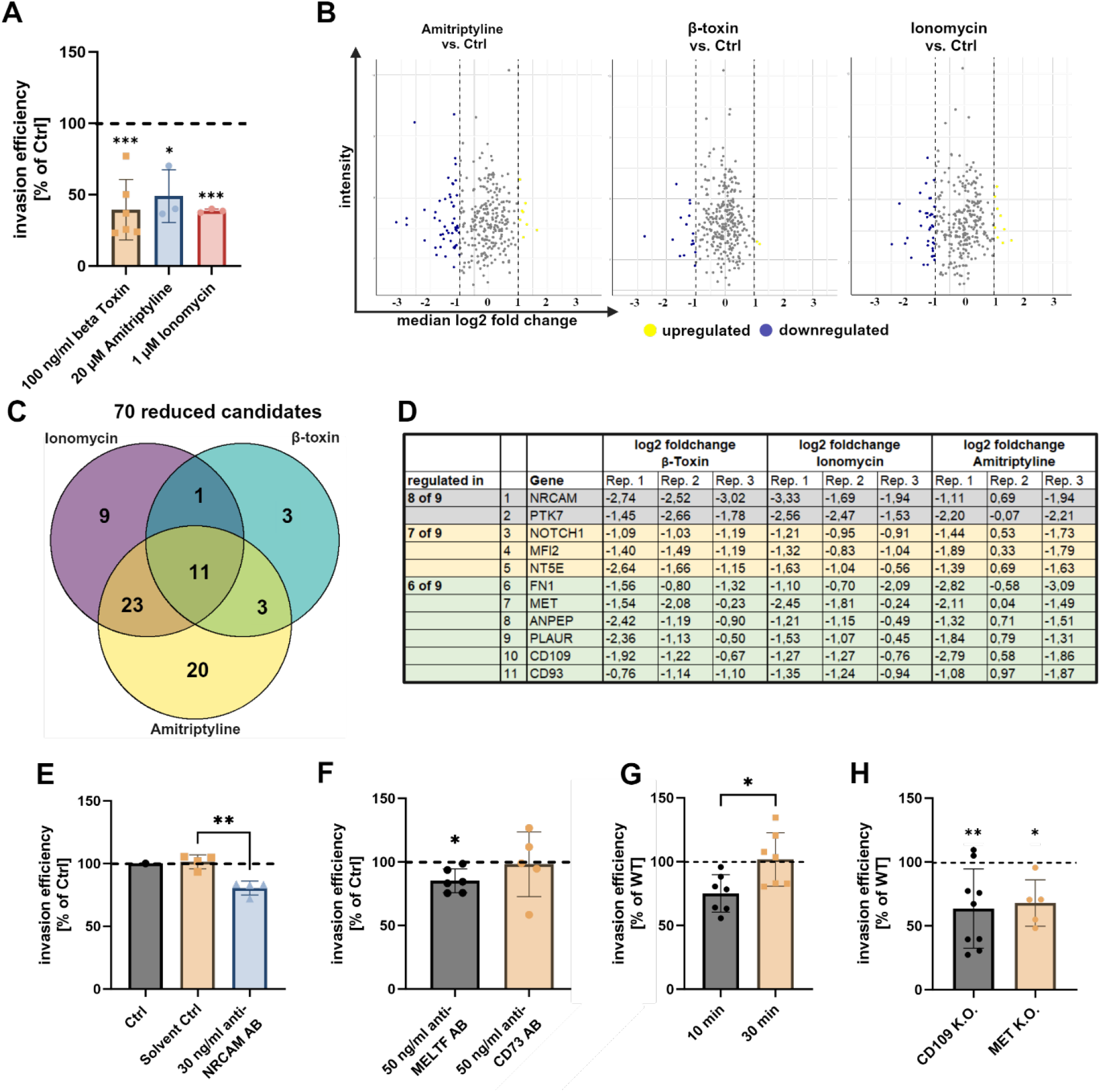
Identification of ASM-dependent host co-receptors for *S. aureus* by an APEX2-based proximity labeling screen. (**A) Decorated bacteria are still internalized by host cells in an ASM-dependent manner**. HuLEC were treated with amitriptyline, ionomycin and β-toxin and then infected with APEX2-decorated *S. aureus* JE2. Invasion efficiency was determined by lysostaphin protection assay and CFU recovery assays. Numbers of bacteria internalized by treated cells were normalized to untreated controls (set to 100%, dotted line). (**B-D) Establishment of the ASM and Ca**^**2+**^**-dependent host-pathogen interactome**. HuLEC were pretreated with either amitriptyline, the bacterial SMase β-toxin, ionomycin or left untreated. Cells were infected with APEX2-decorated *S. aureus* and interaction partners were determined by proximity labeling. (B) Abundance of individual proteins was compared to untreated controls and median log2 fold changes were determined (n=3). (C) Proteins with a median log2 fold change ≤ -1 were selected as candidate targets for each condition. (D) List of candidate receptors that were downregulated in at least 6 of 9 replicates (regardless of the treatment) were considered as „consistently reduced”. **(E, F) Blockade of NRCAM and MELTF with antibodies reduces invasion efficiency of *S. aureus***. HuLEC cells were pretreated with antibodies targeting NRCAM (G) or melanotransferrin (MELTF), NT5E/CD73 (H) or the respective solvent control. Then, cells were infected with *S. aureus* for 10 min and invasion efficiency was determined by a lysostaphin protection assay and CFU counting (n≥4). **(G) *S. aureus* invasion is reduced in a PTK7 K.O. cell line**. Invasion efficiency of *S. aureus* JE2 after 10 min or 30 min was determined in a HeLa cell line lacking PTK7 and was compared to WT cells (n=7). **(H) *S. aureus* invasion is reduced in a CD109 and MET K.O. cell lines**. Invasion efficiency of *S. aureus* JE2 after 10 min was determined in a HeLa cell line lacking CD109 (n=9) and MET (n=5) was compared to WT cells. Statistics: unpaired Student’s t-test (E, G) or one sample t-test (F, H). *p≤0.05, **p≤0.01.

Mass spectrometric comparison of treated samples to the untreated control (**Figure 2**, B, **Extended Data 2**) revealed eleven proteins which were found decreased in their abundance in all treatment conditions (**Figure 2**, C, D).

These included neuronal adhesion molecule (NRCAM), protein tyrosine kinase 7 (PTK7), 5’-nucleotidase CD73 (NT5E), melanotransferrin (MELTF), neurogenic locus notch homolog protein 1 (NOTCH1), the protein-tyrosine kinase Met and CD109.

To validate our findings, we treated HuLEC with an anti-NRCAM antibody prior to infection. This reduced rapid *S. aureus* invasion by ∼20% when compared to the control (**Figure 2**, E). While antibody blocking of MELTF also slightly affected invasion, an antibody directed against CD73 failed to do so (**Figure 2**, F). A HeLa PTK7 K.O. cell pool (**Supp. Figure 1**, B) showed decreased *S. aureus* invasion for short (10 min) but not for longer (30 min) infection times, suggesting an involvement of PTK7 in the rapid Ca^2+^-/ASM-dependent internalization of *S. aureus* (**Figure 2**, G). Similarly, we observed a decreased internalization of *S. aureus* in cells lacking CD109 or the tyrosine-protein kinase Met at 10 min p.i. (**Figure 2**, H, **Supp. Figure 2**, C).

### Filtering for validated surface proteins increases the hit rate for *S. aureus* receptors

We next selected proteins among the 305 candidates that are validated surface proteins according to a cell surface protein atlas [41]. We identified 89 proteins that were detected by proximity labeling and possess validated host surface localization from which many are known interaction partners of pathogens including *S. aureus* (**Figure 3**, A, see **Extended Data 1** for corresponding references). This analysis includes receptors validated in the present study by antibody blocking or genetic ablation. Selecting for host cell surface proteins increased the hit rate for known interaction partners of pathogens/ *S. aureus* (**Figure 3**, B and C; **Extended Data 1**). The unfiltered data set contained 22% of known pathogen interaction partners (8% for *S. aureus*, 14% for other pathogens; **Figure 3**, B). This proportion increased to 53% after selecting for validated surface proteins (20% for *S. aureus*, 33% for other pathogens; **Figure 3**, C).

**Figure 3.**
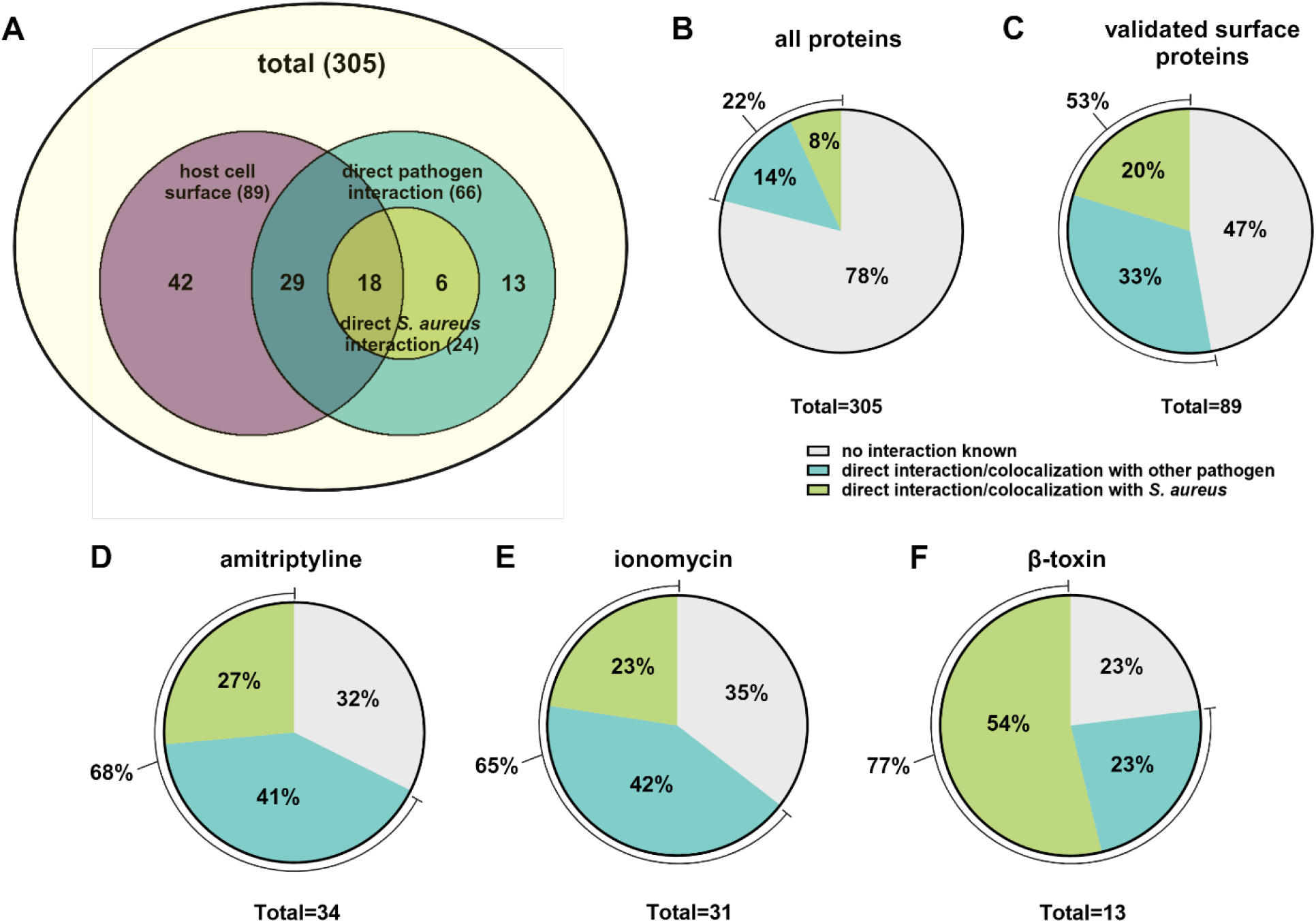
Enrichment of proteins with known pathogen interaction by selecting validated surface proteins. 89 of the 305 *S. aureus* interaction partner candidates identified in the APEX2 screen were selected as validated surface proteins [41]. The data set was then analyzed for proteins that were known to directly interact/localize in close proximity with pathogens (66) or *S. aureus* (24, A) including receptors validated in this study. The proportion of proteins known to interact/colocalize with pathogens or *S. aureus* among all candidates (B) or candidates validated as surface protein (C) was determined. Validated surface proteins that were decreased upon amitriptyline (D), ionomycin (E) or β-toxin (F) treated samples during proximity labeling were analyzed for colocalization/interactions with pathogens or *S. aureus* (including proteins validated in the present study).

This proportion was even higher for candidate receptor proteins that were decreased upon blocking the rapid internalization pathway by amitriptyline (**Figure 3**, D), ionomycin (**Figure 3**, E) or β-toxin **Figure 3**, F). In all three conditions, proximity labeling detected predominantly proteins with already known pathogen interaction.

## Discussion

Identification of receptors by genetic approaches such as RNAi or CRISPR [27] is restricted to receptors that reduce the cellular uptake of the pathogen of interest, while other proteins that are not directly involved in internalization, but might have other function during infection, are missed. Moreover, the reduced invasion of a pathogen after ablation of a host gene, does not necessarily identify the respective gene as host cell receptor, as it may serve other cellular functions important for pathogen uptake such as regulation of membrane composition or actin organization. Cross-linking approaches also enable the identification of receptors [28] but they require molecular interaction between host receptors and pathogen surface. Thus, receptors that are not directly bound to the pathogen surface but might be recruited to pathogen entry site as a co-receptor, are not detected. Additionally, analysis of cross-linked proteins from complex samples, such as whole cell lysates, is challenging [42].

We here present a novel technique for the identification of receptors, which is based on proximity biotinylation. Compared to other approaches that previously were used to identify host cell receptors, the APEX2-based method has important advantages. Identification of proteins by proximity labeling requires close vicinity to the pathogen but does not need an interaction at molecular level. Moreover, proximity biotinylation is not restricted to receptors that are directly involved in the internalization of the pathogen. Hence, we think that the APEX2-based approach enables the detection of host proteins that are present at host-pathogen contact sites in their entirety. However, APEX2-based biotinylation also detects bystander proteins, which are randomly localized at the site of infection.

We hypothesized that a specific set of host cell surface receptors is required for the rapid uptake of *S. aureus* by endothelial cells [26]. Since the identification of receptors that are involved in fast invasion processes is exceptionally demanding, we designed an assay for proximity labeling of host surface receptors using APEX2-decorated bacteria (**Figure 1**, A). Compared to other techniques such as BioID and TurboID [30, 31], APEX2 allows for proximity biotinylation with high temporal resolution [29]. Our approach identified 305 candidate proteins (**Extended Data 1**), of which many were shown to interact with *S. aureus* such as α_5_β_1_ integrins [10], fibronectin [36], CD44 [18], annexin 2 [19], Hsp60 [20], Hsc70 [21], Hsp90 [22], Desmoglein 1 [23], corneodesmosin [24], laminin [37] or Ephrin type A receptor 2 [25] and others. Of these candidates we stringently selected proteins (**Figure 2**, D) that were found reduced in conditions that block the rapid ASM-dependent internalization pathway.

Of the resulting eleven proteins we validated PTK7, NRCAM, MELTF, CD109 and MET as novel (co)receptors for ASM-dependent host cell entry of *S. aureus* (**Figure 2**, E-H). PTK7, an inactive tyrosine kinase, was previously linked to extracellular matrix rearrangement, cytoskeleton and focal adhesions [43, 44] and was required exclusively for the early internalization of *S. aureus* (**Figure 2**, G). NRCAM possesses fibronectin-like as well as immunoglobulin-like domains [45], that could interact with staphylococcal fibronectin-binding proteins or protein A, respectively. The tyrosine kinase Met, which we found involved in *S. aureus* invasion, is a well-known host cell receptor of *Listeria monocytogenes* [9]. Moreover, the *Drosophila melanogaster* homolog of CD109, TepIII, was previously shown to play a role in phagocytosis of *S. aureus* [34].

We here suggest that next to primary receptors such as α_5_β_1_ integrins or CD44, which initially interact with the pathogen [10, 18], PTK7, NRCAM, MELTF, CD109 and Met are accessory receptors, whose interaction with the bacteria modifies and accelerates the internalization. Hence, blockade of each receptor had a moderate yet significant impact on *S. aureus* internalization (**Figure 2**, E-H). *S. aureus-* mediated release of ASM by lysosomal exocytosis may facilitate the recruitment of NRCAM, PTK7 and MELTF or other receptors, e.g. via formation of ceramide-enriched platforms. Supporting this model, blocking ASM-dependent internalization did not reduce the interaction of *S. aureus* with known primary receptors such as α_5_β_1_ integrin or CD44.

ASM-dependence was demonstrated for the host cell entry of a variety of pathogens. Hence, we suggest that the surface molecules identified by APEX2-based proximity labeling, represent interesting candidate receptors for other bacteria and viruses. For instance, Met and ANPEP, which also were identified in the screen, are known entry factors for *Listeria* [9] and human corona virus [46], respectively. Interestingly, a rapid ASM-dependent uptake previously was demonstrated for adenovirus [47], suggesting that multiple pathogens use the presented strategy.

In our study, the CWT domain enabled attachment of APEX2 to *S. aureus*. Since the interaction between CWT and the staphylococcal cell wall is highly specific, decoration of other pathogens with APEX2 would require alternative proteins, such as APEX2-conjugated antibodies or bacteriophage receptor-binding proteins [48].

Proximity labelling is prone for detection of false-positive candidates either due to biotinylation of high abundant by-stander proteins or unspecific binding during streptavidin pull down. Nevertheless, we reached a hit rate of ∼20% known *S. aureus* receptors after filtering for surface resident proteins (**Figure 3**, C). Hence, proteins detected by our proximity labeling approach has identified additional potential *S. aureus* surface receptors, which especially is supported by the high proportion of proteins known to interact with other pathogens. Our dataset thus indicates that diverse pathogens use a similar subset of molecules to interact with host cells. It will be interesting to compare host interactomes of different pathogens in future studies.

## Acknowledgements

This work was supported by the Deutsche Forschungsgemeinschaft (DFG; https://www.dfg.de) within the research training group RTG2581 (M.R., F.S.). M.R. was supported by funds of the Bavarian State Ministry of Science and the Arts and the University of Würzburg to the Graduate School of Life Sciences (GSLS), University of Würzburg. The DFG funded the Leica TCS SP5 CLSM under project code 116162193 and the BD FACSAriaIII cell sorter under project code 206080318.

We thank Sibylle Schneider-Schaulies and Thomas Rudel for valuable discussions and critically reading the manuscript and Marie Schöl for help with the biotin pull down. We are indebted to Andreas Schlosser and Stephanie Lamer (Robert-Virchow Zentrum, Würzburg) for mass spectrometry. We thank Julian Fink and Jürgen Seibel for providing the clickable ceramide analog. Figures were created in BioRender.

## Author Contributions

M.R., M.F. conceptualized the work, designed the methodology and wrote the original draft manuscript. M.F. acquired funding. M.R. F.S., A.K., and K.P. performed the experiments; M.R., M.F. supervised the experiments and analyzed the data; all authors edited and revised the manuscript.

## Materials and Methods

### Cell culture

HeLa (ATCC CCL-2™) were cultivated in RPMI+GlutaMAX™ medium (Gibco™, Cat. No. 72400054) supplemented with 10% (v/v) heat-inactivated (56°C at 30 min) fetal bovine serum (FBS, Sigma Aldrich, Cat. No. F7524) and 1 mM sodium pyruvate (Gibco™, Cat.No. 11360088).

HuLEC-5a (ATCC CRL-3244™) were cultured in MCDB131 medium (Gibco™, Cat. No. 10372019) complemented with microvascular growth supplement (Gibco™, Cat. No. S00525), 2 mM GlutaMAX™ (Gibco™, 35050061), 5 % (v/v) heat-inactivated (56°C at 30 min) FBS, 2.76 µM hydrocortisone (Sigma Aldrich, Cat. No. H0888), 0.01 ng/ml human epidermal growth factor (Pep Rotech, AF-100-15) and 1x Penicillin-Streptomycin (Gibco™, Cat. No. 15140122).

HuLEC were detached by StemPro™ Accutase™ (Gibco™, Cat. No. A1110501) and seeded at the indicated density two days prior to the experiment, whereas HeLa were detached with TrypLE™ (Gibco™, Cat. No. 12604013) and seeded one day prior to the experiment.

### Generation of HeLa KO cell lines

Generation of K.O. cell lines is based on a previous protocol [49]. sgRNAs (**Table 1**) were designed with the CHOPCHOP online tool (chopchop.cdu.uib.no) and synthesized as primer pairs (Sigma Aldrich). After phosphorylation with T4 nucleotide kinase (ThermoFisher, Cat. No. ER0031), the sgRNAs were cloned into the BbsI sites of pSpCas9(BB)-p2A-GFP (AddGene #48138, [49]) via forced ligation by BbsI (ThermoFisher, Cat. No. ER1012) and T4 Nulceotide ligase (ThermoFisher, Cat. No. EL0011). The constructs were then transformed in *E. coli* DH5α (ThermoFisher, Cat. No EC0112) and subsequently sequence verified.

**Table 1.**
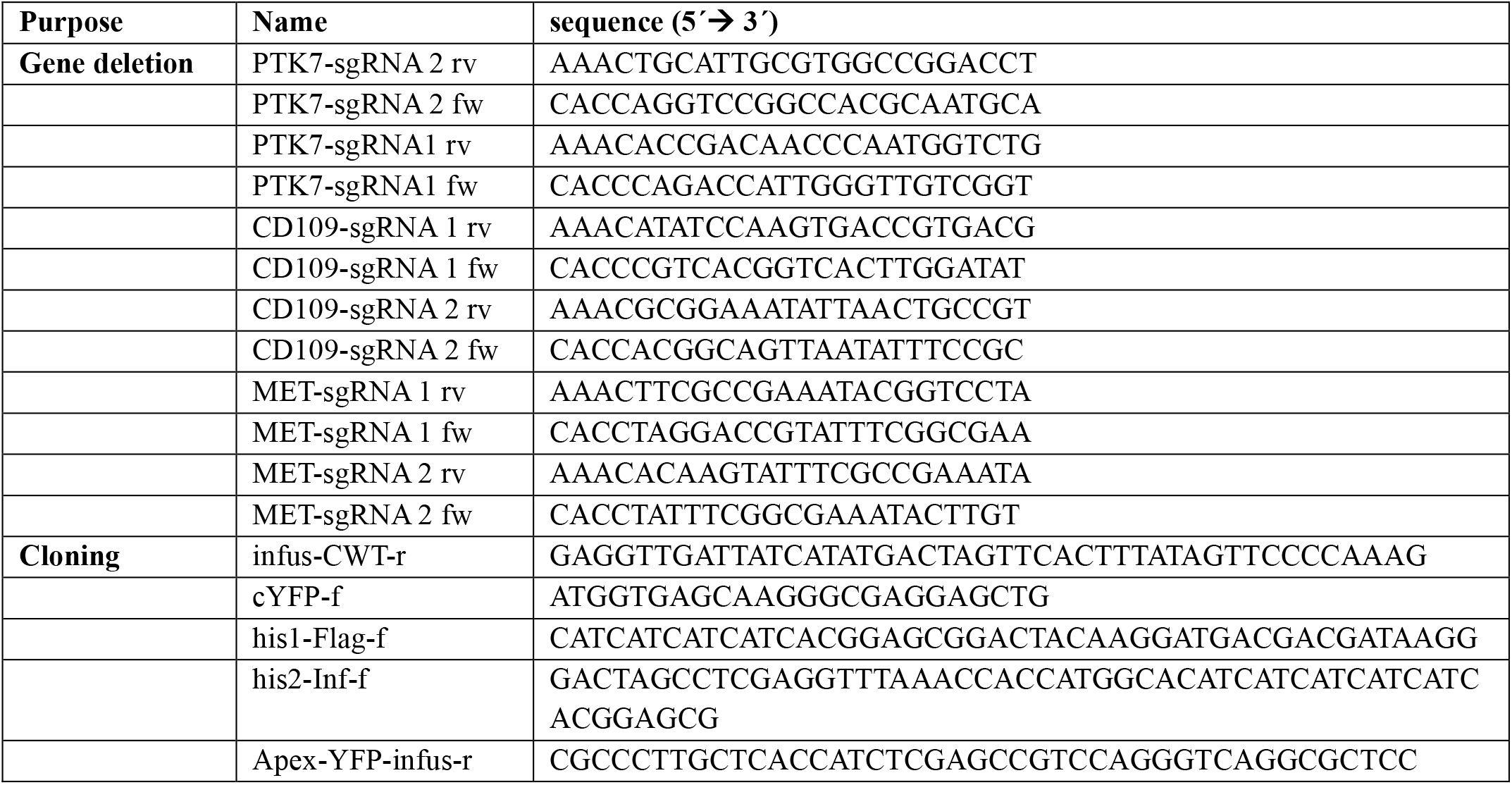
Oligonucleotides used in this study.

**Table 2.** Host cell treatment with various compounds.

| compound | Supplier/Cat.No. | preincubation time | removal prior to infection? |
| --- | --- | --- | --- |
| 30 ng/ml anti-NRCAM antibody | proteintech/<br>21608-1-AP | 75 min | removed prior to infection |
| 50 ng/ml anti-CD73 antibody | Thermo Fisher/<br>41-0200 | 75 min | removed prior to infection |
| 50 ng/ml anti-MELTF antibody | Biozol/BLD-363101 | 75 min | removed prior to infection |
| amitriptyline | Sigma Aldrich/<br>A8404 | 75 min | present during infection |
| Ionomycin | Sigma Aldrich/ I0634 | 75 min | present during infection |
| β-toxin | Sigma Aldrich/<br>S8633 | 75 min | removed prior to infection |

HeLa cells were transfected with 1 μg each of two distinct sgRNA constructs using JetPrime (Polyplus) following the manufacturer’s instructions. After 36-48h, cells were sorted for strong GFP expression in a FACS Aria III (BD Bioscience). The resulting cell pools were cultivated and tested for loss of expression of the respective sgRNA target by Western Blotting.

Following antibodies were used: mouse anti-PTK7 monoclonal IgG1 (bio-techne, Cat. No. MAB4499) with a horse radish peroxidase (HRP)-conjugated anti-mouse antibody (SantaCruz, Cat. No. sc-516102) and rabbit anti-TPC1 polyclonal (Thermo Fisher, Cat. No. PA5-41048) with an HRP-conjugated anti-rabbit antibody (Biozol, Cat. No. 111-035-144).

### Generation of APEX2 construct

*pLV-APEX2-YFP-CWT*. YFP-CWT was amplified from pLVTHM-YFP-CWT [50] with oligonucleotides cYFP-f and infus-CWT-r, thereby introducing overhangs suitable for InFusion cloning. Next we amplified FLAG-APEX from pLX304-Flag-APEX2-NES [51] with his1-Flag-f and APEX-YFP-infus-r, a subsequent reamplification with his2-Inf-f and APEX-YFP-infus-r thereby introducing a N-terminal 6x His-tag. Subsequently, we performed an InFusion reaction with PmeI/SpeI-opened pLVTHM vector [52] as well as 6xHis-FLAG-APEX2 (886 bp) and YFP-CWT (1089 bp) The resulting vector was termed pLV-6xHis-FLAG-APEX2-YFP-CWT.

Constructs were transformed into *E. coli* DH5α and sequences were validated. Plasmid preparations were performed using standard laboratory procedures. VSV-G-pseudotyped Lentiviral particles were generated by transfecting with each lentiviral vector as well as the plasmids pMD2.G (VSV G) and psPAX2 following the protocol of [52] and target cells were transduced as described previously [50].

#### *S. aureus* culture

*S. aureus* liquid cultures were either grown in 37 g/L brain heart infusion (BHI) medium (Thermo Fisher, Cat. No. 237300). *E. coli* liquid cultures were grown in LB medium [10 g/l tryptone/peptone (Roth, Cat. No. 8952.4,) 5g/l yeast extract (Roth, Cat. No 2363.2), 10 g/l NaCl (Sigma Aldrich, Cat. No. S5886)]. For agar plates, 15 g/L agar (Otto Norwald, Cat. No. 257353), was added to either TSB (*S. aureus*) or LB (E. coli).

Media and plates were supplemented with appropriate antibiotics [5 µg/ml erythromycin (Sigma Aldrich, Cat. No. E5389), 10 µg/ml chloramphenicol (Roth, Cat. No. 3886.2), 100 ng/ml Carbenicillin (Roth, Cat. No. 6344.2)].

#### *S. aureus* infections

Host cells were seeded in 6 well plates (2×10^5^ cells per well), 12 well plates (1×10^5^ cells per well) or 24 well plates (0.5×10^5^ cell per well) either one day (HeLa, EA.hy 926, 16HBE14o^-^) or two days (HuLEC, HuVEC) prior to the experiment. Host cells were washed thrice with DPBS and infection medium or Ca^2+^-free infection medium [DMEM w/o calcium (Gibco™, Cat. No. 21068028) complemented with 10 % (v/v) heat-inactivated (56°C at 30 min) FBS and 200 µM BAPTA (Merck Millipore, 196418)] with or without 1.8 mM CaCl_2_ (Roth, Cat. No. 5239.1) was added. If indicated, cells were pretreated with compounds prior to infection (for details about treatments see **Table 1**). For experiments involving blocking of receptors by antibodies, respective solvent controls were implemented (final concentrations in infection medium: 0.002% (w/v) NaN_3_ (Sigma Aldrich, Cat. No. S2002) 5% glycerol (Roth, Cat. No 3783.2) in 10% DPBS for 30 ng/ml anti-NRCAM antibody as well as well as 0.01 % (w/v) NaN_3_ for 50 ng/ml anti-CD73 and 50 ng/ml anti-MELTF antibodies.

For all infections, the *S. aureus* strain JE2 was used [53]. An *S. aureus* overnight culture grown in BHI containing the appropriate antibiotics was diluted to an OD_600_=0.4 in the same medium. The culture was grown to an OD_600_= 0.6-1.0 and 1 ml bacterial suspension was harvested by centrifugation and washed twice with Dulbecco’s Phosphate Buffered Saline (DPBS, Gibco™, Cat. No 14190169). The bacteria were resuspended in infection medium. The number of bacteria per ml in the suspension was determined with a Thoma counting chamber and the MOI was determined. If not indicated otherwise, an MOI=10 was used for infections.

The infection was synchronized by centrifugation 800xg/8 min/RT (end of the centrifugation: t=0). To determine bacterial invasion, the infection was stopped after 30 min (unless indicated otherwise) by removing extracellular bacteria with 20 µg/ml Lysostaphin (AMBI, Cat. No. AMBICINL) in infection medium for 30 min. Then, host cells were washed thrice with DPBS, lysed by addition of 1 ml/well (12 well plate) or 0.5 ml/well (24 well plate) Millipore water and the number of bacteria in lysates was determined by plating serial dilutions (10^−1^, 10^−2^, 10^−3^) on TSB agar plates. Plates were incubated overnight at 37 °C and CFU were enumerated. To determine invasion efficiency, the number of bacteria determined in tested samples were normalized to untreated controls (set to 100%).

### Decoration of *S. aureus* with APEX2-YFP-CWT

HeLa cells expressing APEX2-YFP-CWT were seeded into twelve TC dishes (Ø150 mm) with a density of 6×10^5^ cells per dish and cultured for 5 days (medium was exchanged after 2-3 days). Isolation of the cytoplasm was based on a previously published protocol [54]. Shortly, cells were washed thrice with DPBS and then, cells were scraped from the substratum in 4 ml cold DPBS per dish. The cell suspension for 3 dishes was pooled and samples were centrifuged at 800xg for 7 min at 4°C. The pellet of each cell pool was resuspended in 1 ml Buffer B [20 mM HEPES pH 7.6 (Roth, Cat. No. 6763.3), 220 mM mannitol (Roth, Cat. No. 4175.1), 70 mM sucrose (Roth, Cat. No. 4621.2), 1 mM EDTA (Roth, Cat. No. 8040.2), 1x protease inhibitor cocktail (Sigma Aldrich, Cat. No. 11873580001)] and cells were incubated for 20 min at 4°C. Subsequently, cells were lysed by a tissue homogenizer and lysates were cleared by centrifugation 10,000xg/10 min/4°C. Then, lysates were centrifuged in a Optima™ MAX-XP ultracentrifuge (Beckman Coulter) at 100,000xg at 4°C for 45 min. The supernatants containing the cytosol of the cell pools were combined and concentrated in an Amicon^®^ centrifuge concentrator tube (3 kDa cut off, Sigma Aldrich, Cat. No. UFC901024) by centrifugation at 4,000xg/4°C for 30-45 min until a volume of 1.5-2.0 ml was reached.

For decoration of *S. aureus* with APEX2-YFP-CWT, a *S. aureus* overnight culture was grown in BHI medium, freshly diluted to OD_600_=0.4 in 10 ml BHI medium and grown until a OD_600_=0.6-1.0 was reached. 3 ml of the culture were harvested by centrifugation, washed twice with DPBS and incubated with the concentrated cytoplasm of HeLa APEX2-YFP-CWT for 30 min at RT in an end-over-end rotator. The decorated bacteria were washed once with DPBS, resuspended in 1 ml Hank’s balanced salt solution (HBSS; Gibco™, Cat. No. 14025-100) and the number of bacteria was determined in a Thoma counting chamber. The bacterial suspension was then either used for invasion studies or proximity labeling. To validate decoration of bacteria, the *S. aureus* suspension was analyzed with an Attune NxT flow cytometer (Thermo Fisher, Attune Cytometric Software v5.2.0) gating for YFP fluorescence (Ex. 488nm/Em. bandpass 530/30 nm).

### Proximity labeling with APEX2-decorated bacteria

Based on a published protocol [55], we seeded HuLEC in 6 well plates with a density of 2.5×10^5^ cells per well (3 wells per sample) two days prior to the experiment. Cells were pretreated with infection medium containing 1 µM ionomycin, 100 ng/ml β-toxin or 20 µM amitriptyline for 75 min. Then, we washed the cells thrice with DPBS and added fresh infection medium containing 1 mM biotin-phenol (Sigma Aldrich, SML2135) as well as amitriptyline and ionomycin. At this stage we omitted β-toxin. Subsequently, cells were infected with APEX2-decorated *S. aureus* JE2 at an MOI=50 by centrifuging the bacteria on the host cells at 800 x g for 8 min at RT. After 15 min, cells were washed thrice to remove non-attached bacteria. Then, 1 ml per well HBSS containing 1 mM biotin phenol was applied and biotinylation was initiated by adding 0.003% (v/v) H_2_O_2_ (Sigma Aldrich, Cat. No. H1009) for 1 min. Controls without H_2_O_2_ addition were perform. Immediately, cells were washed once with cold DPBS and 2 ml per well quencher solution [10 mM sodium ascorbate (Sigma Aldrich, Cat. No. A4034), 5 mM Trolox (Sigma Aldrich, Cat. No. 238813), 10 mM sodium azide (Sigma Aldrich, Cat. No. S2002)] were applied to stop biotinylation. Then, cells were either fixed with 0.2 % glutaraldehyde (Sigma Aldrich, Cat. No. 10333) / 4% paraformaldehyde in PBS (Morphisto, Cat. No 11762.01000) for 30 min RT and further prepared for microscopy or processed for streptavidin pull down. Therefore, cells were detached with a cell scraper and three wells per condition were pooled. Cells were centrifuged at 3500 x g for 7 min at 4°C and were resuspended in 100 µL RIPA buffer [50 mM TRIS-HCl (Sigma Aldrich, Cat. No. T1503), 150 mM NaCl (VWR, Cat. No. 27810.364), 1 % (v/v) Triton-X 100 (Roth, Cat. No. 3051.4), 0.1 (w/v) sodium dodecyl sulfate (SDS, Roth, Cat. No. CN30.3), 2.5 (w/v) sodium deoxycholate (Roth, Cat. No. 3484.3), 1x protease inhibitor cocktail (Sigma Aldrich, Cat. No. 11873580001)]. After lysis for 30 min at 4°C, samples were centrifuged 14,000 x g for 10 min at 4°C. Lysates were loaded to an Amico^®^ centrifuge concentrator (3 kDa cut off, Sigma Aldrich, Cat. No. UFC901024) and centrifuged at 4,000 x g for 30 min at 4°C to remove residual biotin phenol until a volume of ∼1 ml was reached. The concentrate was diluted with 5 ml DPBS containing 1x protease inhibitor (Sigma Aldrich, Cat. No. 11873580001) and re-concentrated by centrifuging at 4,000 x g for 30-45 min at 4°C until a volume of 500-100 µL was reached. The same procedure was repeated with 5 ml RIPA buffer. Next, 40 µL per sample of magnetic streptavidin beads (Thermo Fisher, Cat. No. 88816) were equilibrated with 1 ml per sample RIPA buffer and incubated with the concentrated biotin phenol-free cell lysates at 4°C overnight with permanent rotation.

Next, the beads were washed with i.) 2x 1 ml RIPA buffer, ii.) 1x 1 M KCl (Roth, Cat. No. 6781.1), iii.) 1x 0.1 M Na_2_CO_3_ (Roth, Cat. No. 8563.1), iv.) 2 M Urea (Roth, Cat. No. 2317.1) in 10 mM TRIS-HCl pH 8 (Sigma Aldrich, Cat. No. T1503), v.) 1x RIPA. Then, beads were resuspended in 50 µL 6x Laemmli [60 mM TRIS-HCl pH 6.8 (Sigma Aldrich, Cat. No. T1503), 12% (w/v) SDS (Roth, Cat. No. CN30.3), 47% (v/v) glycerol (Roth, Cat. No. 3783.2), dithiothreitol (DTT; Roth, Cat. No. 4227), 0.01% (w/v) bromophenol blue (Sigma Aldrich, Cat. No. B0126)] containing 2 mM biotin (Thermo Fisher, Cat. No 29129) and incubated at 98°C for 10 min to eluate biotinylated proteins. The eluates were analyzed via mass spectrometry.

To verify pull down of biotinylated proteins, the cell lysates of samples before incubation with streptavidin beads (input) were compared with pull down fractions by Western blot. Therefore, 50 µL input samples were mixed with 10 µL 6x Laemmli, boiled at 98°C/10 min and subsequently, analyzed by semi-dry Western blot. PVDF membranes were blocked by 5% milk powder (AppliChem, Cat. No. A0830,5000) in TBS-T pH 7.5 [50 mM TRIS (Sigma Aldrich, Cat No. T1503), 150 mM NaCl (VWR, Cat. No. 27810.364), 0.05% (v/v) Tween^®^ (Roth, Cat. No. 9127.2)]. Membranes were incubated with a horse radish peroxidase (HRP)-linked anti-biotin antibody (Cell Signaling, Cat. No. 7076S) overnight and developed with an HR-16-3200 chemiluminescence reader (Intas).

### Mass spectrometry

#### Single-pot, solid-phase-enhanced sample preparation (SP3)

Samples were processed using an adapted SP3 protocol [56]. Briefly, 200µl reconstitution solution was added to each sample prepared in 50µl Laemmli buffer (Life Technologies), resulting in a DTT concentration of 12mM. Alkylation was performed with 35mM iodoacetamide. 10mM additional DTT was used for quenching. Equal volumes of two types of Sera-Mag Speed Beads (Cytiva, #45152101010250 and #65152105050250) were combined, washed with water and 10 µL of the bead mix were added to each sample. 260µl 100% ethanol was added and samples were incubated for 5min at 24°C, 1000rpm. Beads were captured on a magnetic rack for 2 min, and the supernatant was removed. Beads were washed twice with 200µl 80% ethanol (Chromasolv, Sigma) and then once with 1000µl 80% ethanol. On-bead digest was performed with 0.25µg Trypsin (Gold, Mass Spectrometry Grade, Promega) and 0.25 µg Lys-C (Wako) in 100µl 100mM ammonium bicarbonate at 37°C overnight. Peptides were desalted using C-18 Stage Tips [57]. Each Stage Tip was prepared with three discs of C-18 Empore SPE Discs (3M) in a 200 µl pipette tip. Peptides were eluted with 60 % acetonitrile in 0.1 % formic acid, dried in a vacuum concentrator (Eppendorf), and stored at -20 °C. Peptides were dissolved in 2 % acetonitrile / 0.1 % formic acid prior to nanoLC-MS/MS analysis.

#### NanoLC-MS/MS analysis

NanoLC-MS/MS analyses were performed on an Orbitrap Fusion (Thermo Scientific) equipped with a PicoView Ion Source (New Objective) and coupled to an EASY-nLC 1000 (Thermo Scientific). Peptides were loaded on a trapping column (2 cm x 150 µm ID, PepSep) and separated on a capillary column (30 cm x 150 µm ID, PepSep) both packed with 1.9 µmC18 ReproSil and separated with a 120-minute linear gradient from 3% to 30% acetonitrile and 0.1% formic acid and a flow rate of 500 nl/min. Both MS and MS/MS scans were acquired in the Orbitrap analyzer with a resolution of 60,000 for MS scans and 30,000 for MS/MS scans. HCD fragmentation with 35 % normalized collision energy was applied. A Top Speed data-dependent MS/MS method with a fixed cycle time of 3 s was used. Dynamic exclusion was applied with a repeat count of 1 and an exclusion duration of 90 s; singly charged precursors were excluded from selection. Minimum signal threshold for precursor selection was set to 50,000. Predictive AGC was used with AGC a target value of 4×10^5^ for MS scans and 5×10^4^ for MS/MS scans. EASY-IC was used for internal calibration.

#### MS data analysis

Raw MS data files were analyzed with MaxQuant version 1.6.2.2 [58]. Database search was performed with Andromeda, which is integrated in the utilized version of MaxQuant. The search was performed against the UniProt Human Reference Proteome database (June 2022, UP000005640, 79684 entries), a ***Staphylococcus aureus*** GenBank database (Accession numbers NC_007793, NC_007792, NC_007791, NC_007790) and a small database containing the APEX2 constructs. Additionally, a database containing common contaminants was used. The search was performed with tryptic cleavage specificity with 3 allowed miscleavages. Protein identification was under control of the false-discovery rate (FDR; <1% FDR on protein and peptide spectrum match (PSM) level). In addition to MaxQuant default settings, the search was performed against following variable modifications: Protein N-terminal acetylation, Gln to pyro-Glu formation (N-term. Gln) and oxidation (Met). Carbamidomethyl (Cys) was set as fixed modification. Further data analysis was performed using R scripts developed in-house. LFQ intensities were used for protein quantitation [59]. Proteins with less than two razor/unique peptides were removed. Missing LFQ intensities were imputed with values close to the baseline. Data imputation was performed with values from a standard normal distribution with a mean of the 5% quantile of the combined log10-transformed LFQ intensities and a standard deviation of 0.1.

For the identification of significantly enriched proteins, median log2 transformed protein ratios were calculated from the three replicate experiments and boxplot outliers were identified in intensity bins of at least 300 proteins. Log2 transformed protein ratios of sample versus control with values outside a 1.5x (significance 1) or 3x (significance 2) interquartile range (IQR), respectively, were considered as significantly enriched in the individual replicates.

### Microscopy

HeLa cells were incubated with 10 µM α-NH_2_-ω-N_3_-C_6_-ceramide [33] for 30 min prior to infection with APEX2-decorated bacteria. After proximity biotinylation, samples were fixed with 0.2 % (v/v) glutaraldehyde (Sigma Aldrich, Cat. No. 10333) / 4% paraformaldehyde in PBS (Morphisto, Cat. No 11762.01000) for 30 min RT. Cells were permeabilized with 0.2% (v/v) Triton X-100 (Thermo Fisher, Cat. No. 28314) for 20 min RT and α-NH_2_-ω-N_3_-C_6_-ceramide was stained with 2 µM BODIPY-FL-DBCO (Jena Biosciences, Cat. No. CLK-040-05) for 30 min at 37°C. Then, samples were blocked with 10 % FBS in PBS for 1h RT. *S. aureus* was stained with 25 ng/ml anti-*S. aureus* antibody (Thermo Fisher, PA1-7246) at 4°C overnight and an AlexaFluor™405-conjugated secondary antibody (Thermo Fisher, Cat. No. A48254) for 1h at RT. Biotin was stained by 5 ng/ml streptavidin-PE (Becton Dickinson, Cat. No. 554061) at 4°C overnight. Imaging was performed at a confocal SP5 TCS microscopy (Wetzlar, Germany; Software Leica LAS AF Version 2.7.3.9723) with a 40x immersion oil objective (NA1.3) and a resolution of 2048×2048 pixels.

### R analysis

Mass spectrometry (MS) results from APEX2 proximity labeling screens were processed in R studio [60]. Candidates that were identified in all three biological replicates were selected for further analysis (in total 354 proteins). First, the targets were categorized by the pathfindR package [61] according to the Kyoto encyclopedia of genes and genomes (KEGG, [62-64]). In the input matrix, fold-changes and p-values were arbitrarily set to 1 and 0.05, respectively. Proteins listed in the category “ribosomes” were removed from the dataset for further processing (**Extended Data 1**). Then, number of candidates in other categories were counted and plotted with ggplot [65].

Log_2_ fold changes (FCs) were calculated based on label-free quantification (LFQ) detected during MS analysis in treated conditions vs. LFQs detected in untreated control infections. The log_2_ FCs were rounded to one decimal place and a threshold of log_2_FC < -1 for down- and log_2_FC > 1 for up-regulated was used to identify candidates of which abundance was changed by the treatments. Candidates that were downregulated in 2 of 3 replicate (median log_2_FC < -1) were selected as consistently reduced targets. Proteins that were consistently reduced in all three treatment conditions were targets involved in the rapid ASM-dependent invasion pathway (see **Figure 2**, D).

### Statistical analysis

Statistical analysis was performed in GraphPad prism (V10.1.2). One-sample t-test was used for analysis of normalized data sets. Otherwise, one- or two-way ANOVA, dependent on the number of variables, was used in combination with suitable multiple comparisons testing. Details about sample size and deployed statistical analysis can be found in respective figure legends. All data are shown as mean ± standard deviation.

## Supplementary information

**Supp. Figure 1.**
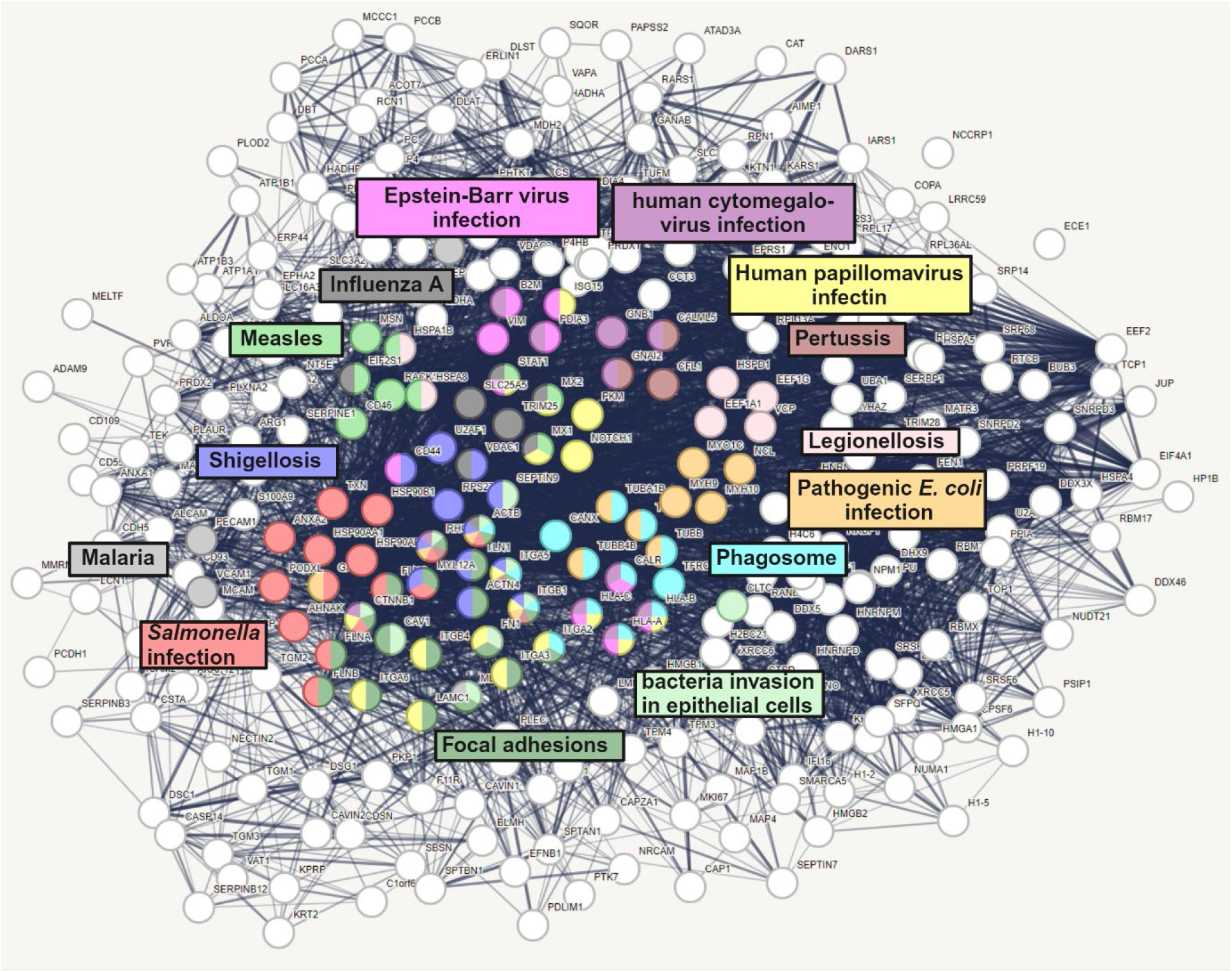
*S. aureus*-host interactome is a highly connected network containing proteins associated with infection. Proteins that were identified by APEX2-based proximity labeling as *S. aureus* interaction partners were analyzed via StringDB. The resulting network has a PPI enrichment p-value < 1×10^−16^. Several proteins that were previously associated with infections for instance, bacteria invasion in epithelial cells (FDR 1×10^−5^), phagosome (FDR 3.75×10^−6^), Shigellosis (FDR 0.0019), Malaria (FDR 0.0308), Salmonella infection (FDR 3.28×10^−5^), pathogenic E.coli infection (FDR 0.00068), Legionellosis (FDR 0.0026), Pertussis (FDR 0.0079), human papillomavirus infection (FDR 9.22×10^−5^), human cytomegalovirus infection (FDR 0.0053), Epstein-Barr virus infection (FDR 0.0072), Influenza A (FDR 0.0227), Measles (FDR 0.0031) as well as focal adhesions (FDR 5.18×10^−8^), the entry site of *S. aureus* into host cells.

**Supp. Figure 2.**
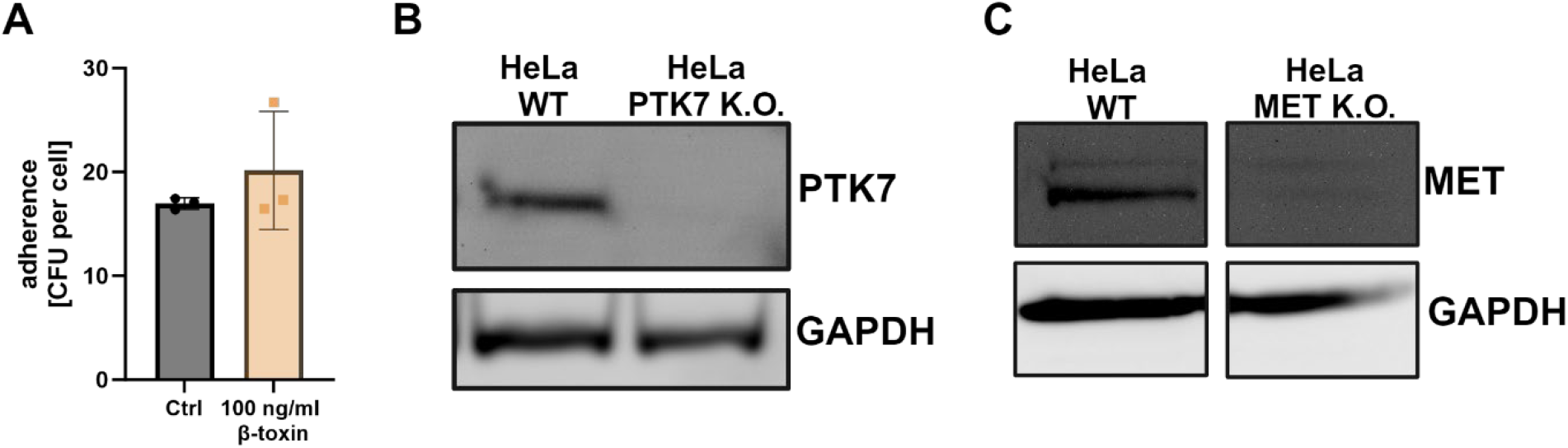
Pretreatment with bacterial sphingomyelinase does not affect the adherence of S. aureus and validation of a PTK7 K.O. cell line. **(A) β-toxin treatment does not affect the adherence of APEX2-decorated bacteria to host cells**. HuLEC were treated with β-toxin and infected with APEX2-decorated bacteria at MOI=50. The number of bacteria that adhered to host cells was determined by CFU plating **(B, C) Western blot demonstrates absence of PTK7 or MET in cas9-treated HeLa cell pool**. HeLa WT cells or a HeLa cell pool lacking PTK7 (B) or MET (C) generated by CRISPR-Cas9 were analyzed for the presence of PTK7 or MET by Western blot.

